# Integrating Lateral Super-resolution and Axial Progression Reveals Distinct Clathrin Pit Formation Pathways

**DOI:** 10.64898/2026.03.06.710132

**Authors:** Cristopher Thompson, Aritra Mondal, Gregory Lafyatis, Comert Kural

## Abstract

Clathrin-mediated endocytosis (CME) relies on the dynamic assembly and remodeling of clathrin coats to drive membrane curvature and vesicle formation at the plasma membrane. Although live-cell fluorescence microscopy has provided critical insights into the timing and molecular composition of endocytic events, directly linking the nanoscale lateral organization of clathrin coats to their three-dimensional progression in real time has remained challenging. Structural approaches such as electron microscopy provide detailed snapshots of clathrin architecture but are inherently static, whereas axial TIRF-based methods report membrane-proximal position with limited lateral resolution. Here, we introduce variable-angle total internal reflection fluorescence structured illumination microscopy (vaTIRF-SIM), a live-cell imaging strategy that integrates lateral super-resolution with dynamic axial sensitivity near the plasma membrane. By combining TIRF-SIM with controlled variation of the evanescent field penetration depth, vaTIRF-SIM enables simultaneous visualization of clathrin coat architecture and relative axial displacement with high spatial and temporal resolution. Applying this approach to de novo clathrin-coated pits reveals coordinated lateral growth and progressive axial advancement from early stages of pit formation through maturation, consistent with early curvature generation that intensifies over time. Extending this analysis to clathrin plaques uncovers two distinct plaque-associated endocytic behaviors: slowly maturing pits that originate at plaque peripheries and progress similarly to de novo pits, and rapid plaque subdomain internalization events marked by accelerated axial progression. Together, these results establish vaTIRF-SIM as an approach that, for the first time, enables direct real-time coupling of nanoscale clathrin coat organization with axial progression during CME in living cells.

## 1 INTRODUCTION

Clathrin-mediated endocytosis (CME) is a fundamental and evolutionarily conserved mechanism by which eukaryotic cells internalize transmembrane receptors, transporters, and associated lipids from the plasma membrane into the endosomal network, thereby maintaining membrane composition and modulating signal transduction (Conner and Schmid 2003; McMahon and Boucrot 2011; Robinson 2015). CME is initiated by the recruitment of adaptor proteins such as the AP-2 complex to phosphatidylinositol-4,5-bisphosphate (PIP2)–enriched membrane domains, followed by the assembly of clathrin triskelions into a lattice that progressively deforms the membrane into a coated pit (Ehrlich et al. 2004; Emanuele Cocucci et al. 2012; Nathan M. Willy, Colombo, et al. 2021). As the coat matures, accessory proteins coordinate cargo selection, membrane curvature generation, and ultimately dynamin-mediated scission to release a clathrin-coated vesicle into the cytosol (Kirchhausen et al. 2014; E Cocucci et al. 2014).

Although CME was long envisioned as a uniform process producing small, rapidly budding coated pits, high-resolution imaging and ultrastructural studies have revealed a broader spectrum of clathrin-coated assemblies at the plasma membrane. In addition to canonical clathrin-coated pits, cells can form larger, flatter, and longer-lived clathrin lattices, commonly referred to as clathrin plaques (Saffarian et al. 2009; Grove et al. 2014). These plaques are particularly prominent at adherent cell surfaces and display distinct dynamics, spatial organization, and dependencies on the actin cytoskeleton compared with classical pits (Traub 2009; Batchelder and Yarar 2010; Lampe et al. 2016). Recognition of this structural and dynamic heterogeneity has refined mechanistic models of CME, emphasizing the plasticity of clathrin coat architecture and the interplay between membrane mechanics, cytoskeletal forces, and endocytic function (N. M. Willy et al. 2017; Ferguson et al. 2017; Umidahan Djakbarova et al. 2021; Umida Djakbarova et al. 2024).

A mechanistic understanding of clathrin coat assembly, maturation, and disassembly has relied heavily on live-cell fluorescence microscopy, which allows real-time tracking of individual endocytic events in living cells (Kural and Kirchhausen 2012; Ferguson et al. 2016; Wu and Kural 2025). Early studies using total internal reflection fluorescence (TIRF) microscopy established characteristic lifetimes of clathrin-coated structures at the plasma membrane, enabling discrimination between productive coated pits, plaques and abortive endocytic events and revealing the staged recruitment of accessory proteins during pit maturation (Saffarian et al. 2009; Loerke et al. 2012; Emanuele Cocucci et al. 2012; Aguet et al. 2013). Multicolor live-cell imaging further allowed direct correlation of clathrin assembly dynamics with cargo engagement and scission timing, while quantitative analysis of fluorescence intensity and temporal profiles provided kinetic measurements describing the timing and rates of individual steps during CME (Taylor et al. 2011). Together, these approaches demonstrated that CME is a highly regulated, multistep process characterized by substantial heterogeneity in both timing and molecular composition.

However, while conventional fluorescence microscopy excels at capturing near-membrane dynamics with high temporal resolution (Mettlen and Danuser 2014), its diffraction-limited spatial resolution constrains direct visualization of the nanometer-scale structural rearrangements within clathrin coats themselves. The lateral diffraction limit of ∼200–300 nm obscures key features of endocytic pit architecture, including membrane curvature, adaptor organization, and lattice remodeling, and can mask the presence of multiple nearby clathrin coats or aggregates that appear as a single structure, limiting our ability to directly link molecular assembly to membrane deformation.

Recent developments in super-resolution fluorescence microscopy have addressed these limitations by achieving nanoscale spatial resolution while maintaining compatibility with live-cell imaging. These methods exceed the diffraction limit of light, allowing for the detailed visualization of cellular structures at or below 100 nm (Gustafsson 2000; Jones et al. 2011). Among these approaches, SIM—and in particular total internal reflection fluorescence SIM (TIRF-SIM)—combines enhanced lateral resolution with selective axial illumination near the plasma membrane, providing high-contrast imaging with reduced phototoxicity and acquisition speeds well suited for rapidly evolving endocytic processes (Li et al. 2015). By resolving individual clathrin coats within dense membrane regions, TIRF-SIM enables separation of isolated *de novo* pits from plaque-associated events and from dynamic aggregates of multiple clathrin-coated structures that are indistinguishable under diffraction-limited imaging (Nathan M. Willy, Ferguson, et al. 2021).

The superior resolution of TIRF-SIM has been instrumental in recent investigations that capture, in real time, the onset of curvature and structural remodeling of clathrin coats in living cells and developing tissues. These findings indicate that curvature is initiated early in pit formation and uncover adaptor clustering previously hidden by the limitations of diffraction-limited imaging (Nathan M. Willy, Ferguson, et al. 2021). By enabling quantitative spatiotemporal analyses of protein organization and membrane morphology at the nanoscale, TIRF-SIM exposes assembly pathways and dynamic intermediates that are invisible to conventional fluorescence microscopy (Akatay et al. 2022). Together, these advances position super-resolution live-cell imaging as a powerful framework for integrating the dynamic, structural, and mechanistic principles governing CME.

Because membrane invagination and vesicle formation during CME involve progressive changes not only in membrane curvature but also in axial position relative to the plasma membrane, capturing the full three-dimensional evolution of clathrin coats presents a fundamental imaging challenge. While conventional TIRF microscopy provides excellent temporal resolution and selective excitation near the cell surface, it is intrinsically limited to two-dimensional readouts. To overcome this limitation, several TIRF-based approaches have been developed that extract axial information while preserving the temporal resolution required to resolve CME dynamics (Saffarian et al. 2009; Nawara and Mattheyses 2023; Wang et al. 2020; He et al. 2025).

Variable-angle TIRF determines the three-dimensional organization of cellular structures by monitoring changes in fluorescence intensity as the angle of incidence is varied. Because the penetration depth of the evanescent field depends on the incident angle, systematic modulation of this angle enables axial positions to be inferred from intensity differences across angles (Burmeister et al. 1994; Cardoso Dos Santos et al. 2016). While this approach provides quantitative sensitivity to axial displacement during endocytic progression, it remains limited by the diffraction-limited lateral resolution of conventional fluorescence microscopy, precluding direct visualization of nanoscale rearrangements within the clathrin coat. Because clathrin assembly, curvature generation, and inward membrane movement are tightly coupled during CME—and because isolated pits, plaque-associated events, and clustered aggregates can coexist within crowded membrane regions—resolving these processes requires simultaneous access to both lateral nanoscale organization and three-dimensional positional information. Here, we combine TIRF-SIM with variable-angle excitation (vaTIRF-SIM) to integrate enhanced lateral resolution with sensitivity to axial progression. This integrated approach enables reliable discrimination of distinct modes of clathrin pit formation, including isolated *de novo* pits, plaque-associated events, and clustered aggregates, while simultaneously tracking their three-dimensional evolution in real time at the plasma membrane of living cells.

By enabling simultaneous nanoscale lateral resolution and dynamic axial readout, vaTIRF-SIM provides, for the first time, a means to directly observe in real time how distinct clathrin architectures generate membrane curvature during CME. This integrated approach makes it possible to distinguish isolated *de novo* clathrin-coated pits—characterized by coordinated lateral growth coupled to axial invagination—from plaque-associated pit formation, in which curvature emerges locally within pre-existing, extended clathrin lattices. In contrast to *de novo* pits, plaque-associated endocytic events frequently exhibit spatially heterogeneous axial progression, the coexistence of multiple invaginating structures within a single lattice, and complex lateral organization that is collapsed or obscured under diffraction-limited imaging. By resolving both the nanoscale lateral arrangement of clathrin coats and their three-dimensional progression as these events unfold, vaTIRF-SIM enables direct, real-time comparison of the spatiotemporal maturation pathways of distinct endocytic modes within the same cellular context.

## 2 RESULTS

### 2.1 vaTIRF-SIM combines lateral super-resolution with real-time axial information

Live-cell studies of clathrin-mediated endocytosis require imaging strategies that can resolve both the nanoscale lateral organization of clathrin coats and their axial progression relative to the plasma membrane. While TIRF-based approaches provide excellent sensitivity to membrane-proximal dynamics and have been widely used to quantify the timing and kinetics of CME, conventional TIRF imaging lacks the lateral resolution necessary to resolve closely spaced pits, plaque-associated structures, and compact aggregates. Conversely, structured illumination microscopy (SIM) improves lateral resolution beyond the diffraction limit and is well suited for live-cell imaging of endocytic structures due to its relatively fast acquisition and moderate phototoxicity, but it does not intrinsically report axial position within the evanescent field.

To directly link nanoscale lateral organization of clathrin coats with their three-dimensional progression during endocytosis, we developed a variable-angle TIRF structured illumination microscopy (vaTIRF-SIM) strategy (Figure 1). This approach integrates the enhanced lateral resolution of TIRF-SIM with axial sensitivity derived from controlled variation of the TIRF incidence angle, while preserving the temporal resolution required to capture clathrin coat dynamics in living cells.

**Figure 1.**
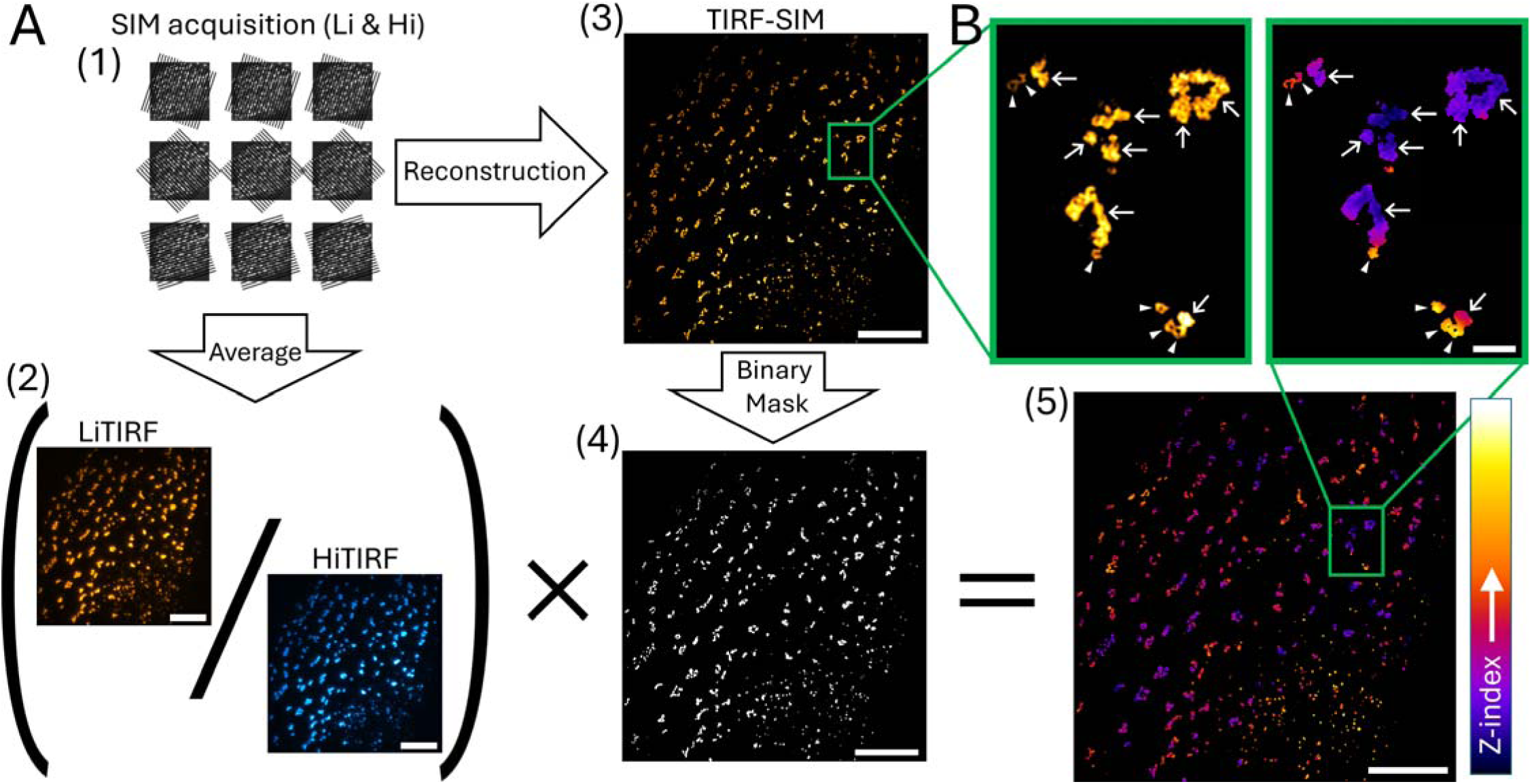
vaTIRF-SIM integrates super-resolved lateral organization with real-time axial information. **A.** The workflow of variable-angle TIRF structured illumination microscopy (vaTIRF-SIM). (1) SIM datasets are acquired sequentially at low- and high-incidence angles (Li and Hi), which produce evanescent fields with deeper and more surface-restricted penetration, respectively (see Methods for optical parameters). (2) Averaged Li and HiTIRF images used to compute axial information in the absence of SIM reconstruction. Pixel-wise division of the averaged Li and HiTIRF images yields a ratiometric axial index that reports relative axial position within the evanescent field but remains diffraction-limited laterally. (3) SIM reconstruction recovers high-spatial-frequency lateral information while preserving the differential axial sensitivity of the two incidence angles. (4) SIM-based segmentation of clathrin-positive structures. The enhanced lateral resolution of TIRF-SIM enables reliable identification and separation of individual clathrin-coated pits, plaque-associated assemblies, and clustered aggregates that cannot be resolved under diffraction-limited imaging. (5) Z-index map computed as the pixel-wise ratio of LiTIRF to HiTIRF fluorescence intensities and masked using the SIM-derived binary maps. Warmer colors indicate higher Z-index values corresponding to greater axial displacement from the membrane–glass interface, whereas cooler colors indicate more surface-proximal localization. Scale bars, 10 µm. **B.** Representative vaTIRF-SIM image illustrating simultaneous visualization of lateral clathrin coat architecture and relative axial progression at the plasma membrane of a SUM-159 cell. Extended, flat clathrin lattices (plaques) are indicated by arrows, whereas highly curved, ring-shaped clathrin-coated pits are marked by arrowheads. The Z-index reveals axial heterogeneity within and between clathrin assemblies, highlighting localized invagination events within crowded membrane regions. Scale bar, 1 µm.

To implement vaTIRF-SIM in living cells, we performed all experiments in genome-edited SUM-159 cells expressing AP2-EGFP from the endogenous locus (Aguet et al. 2016), enabling quantitative visualization of clathrin-mediated endocytic events under physiological expression levels. As illustrated in Figure 1A, SIM datasets were acquired sequentially at two TIRF incidence angles: a low-incidence angle (Li), which produces a deeper evanescent field penetration, and a high-incidence angle (Hi), which restricts excitation to a more surface-proximal region (see Methods for optical parameters). To extract axial information independently of lateral super-resolution, Li and HiTIRF images were first averaged across SIM phase and orientation angles and used to compute a pixel-wise axial index defined as the ratio of LiTIRF to HiTIRF fluorescence intensities (Figure 1A, 1&2). This ratiometric map reports relative axial position within the evanescent field but remains diffraction-limited laterally.

**Figure 2.**
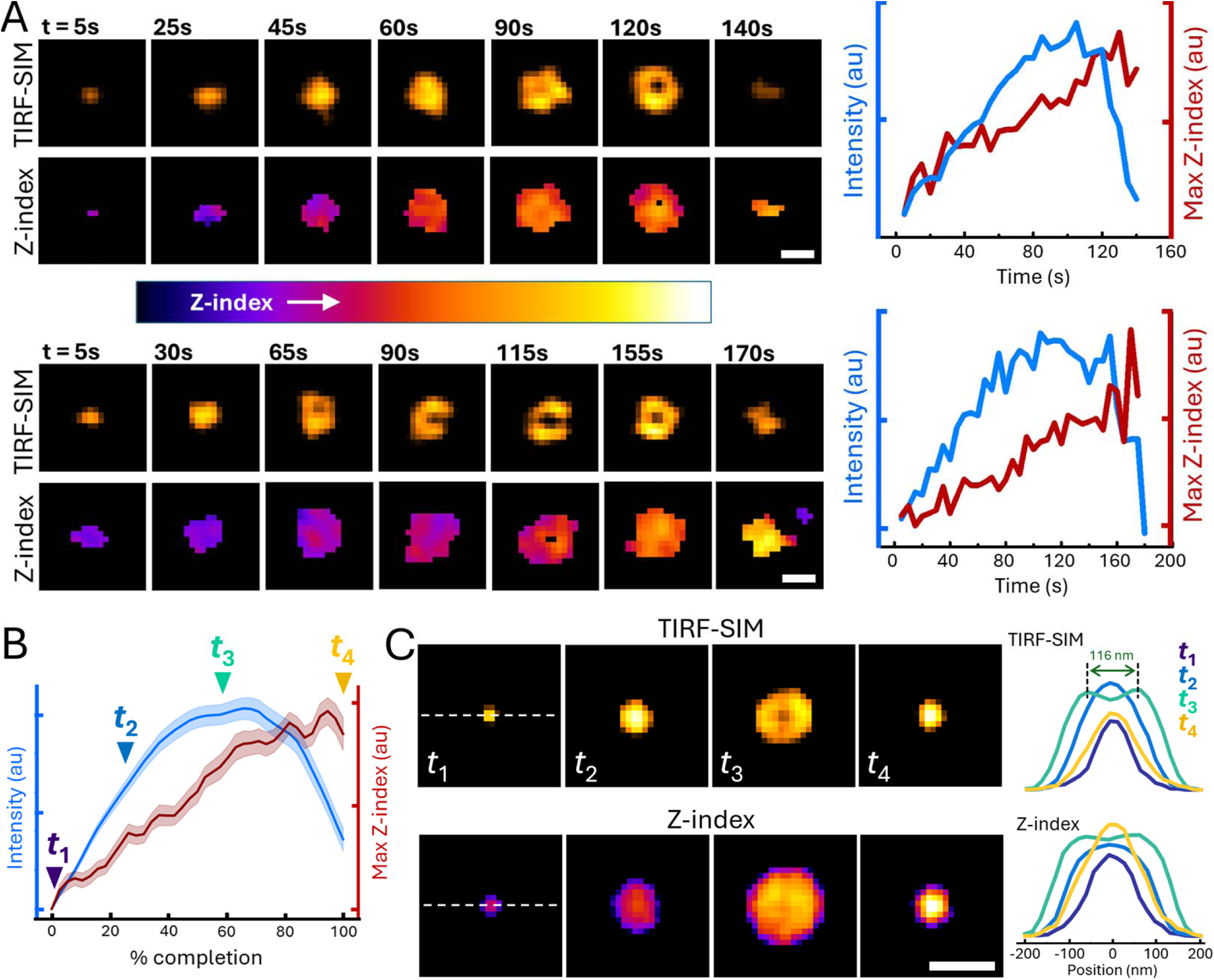
vaTIRF-SIM reveals coordinated growth and axial progression during de novo clathrin pit formation. **A.** Representative examples of *de novo* clathrin pits imaged by vaTIRF-SIM. Left panels show TIRF-SIM images revealing nanoscale lateral organization of clathrin coats over time, while corresponding Z-index maps report relative axial displacement from the plasma membrane. *De novo* pits evolve from compact assemblies to structures with a ring-like organization prior to disappearance. Right panels show time traces of AP2 fluorescence intensity (blue) and maximum Z-index (red) measured within the segmented pit region, illustrating coordinated lateral assembly and progressive axial progression during pit maturation. **B.** Population-averaged dynamics of *de novo* pit maturation. Mean fluorescence intensity (blue) and mean maximum Z-index (red) are shown as a function of normalized lifetime (0–100%), computed from 75 *de novo* pits. Shaded regions indicate ± SEM. **C.** Average images corresponding to the population in (B), computed after lifetime normalization and spatial alignment of individual pits by their centroid (Movie 1). TIRF-SIM images (top) and Z-index maps (bottom) are shown at four representative time points (*t*lJ–*t*lJ). Right panels show line profiles taken at the indicated time points, revealing progressive changes in lateral organization and axial distribution during pit maturation. Scale bars, 0.2 µm.

In parallel, the full SIM dataset was reconstructed independently to recover high-spatial-frequency lateral information (Figure 1A, 3). The resulting TIRF-SIM images resolve individual clathrin-coated pits, plaque-associated assemblies, and clustered aggregates that are not separable under conventional TIRF illumination. These SIM reconstructions were used to segment clathrin-positive structures with high lateral fidelity, generating binary masks that define the spatial footprint of clathrin signal (Figure 1A, 4).

**Figure 3.**
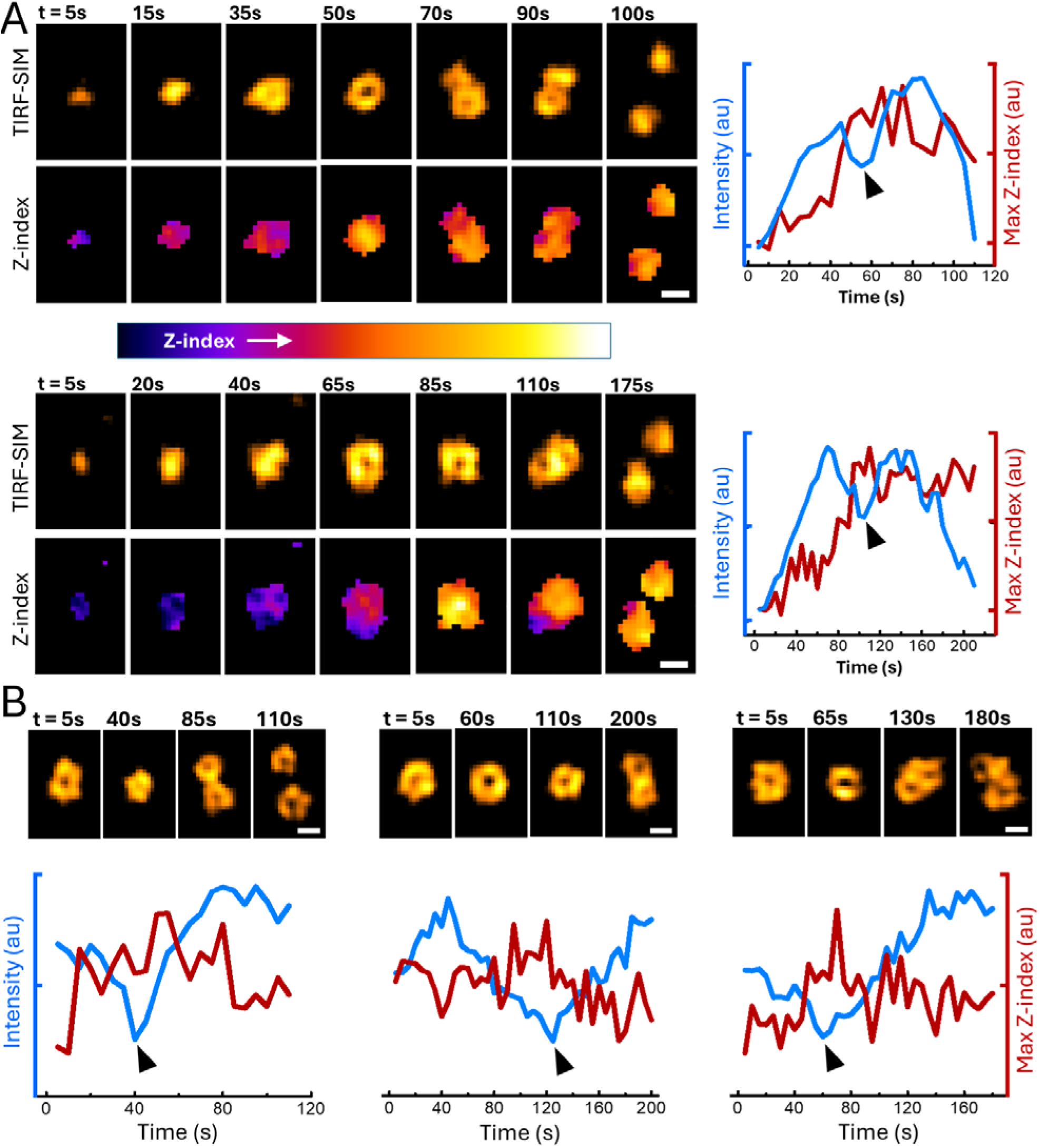
Late-stage splitting of clathrin-coated pits. **A.** Representative *de novo* clathrin pits imaged by vaTIRF-SIM over their full lifetime, from initial detection through coat assembly, axial progression, ring formation, and subsequent splitting. TIRF-SIM images (top rows) resolve nanoscale lateral organization of clathrin coats, while corresponding Z-index maps report relative axial position with respect to the plasma membrane. Right panels show time traces of AP2 fluorescence intensity (blue) and maximum Z-index (red). **B.** Additional examples focusing specifically on mature clathrin pits, with traces beginning after the characteristic ring pattern is already apparent. Across both (A) and (B), splitting of ring-patterned pits is consistently preceded by a pronounced but transient decrease in AP2 fluorescence intensity (arrowheads), while Z-index values remain elevated or fluctuate at high levels. Scale bars, 0.2 µm.

**Figure 4.**
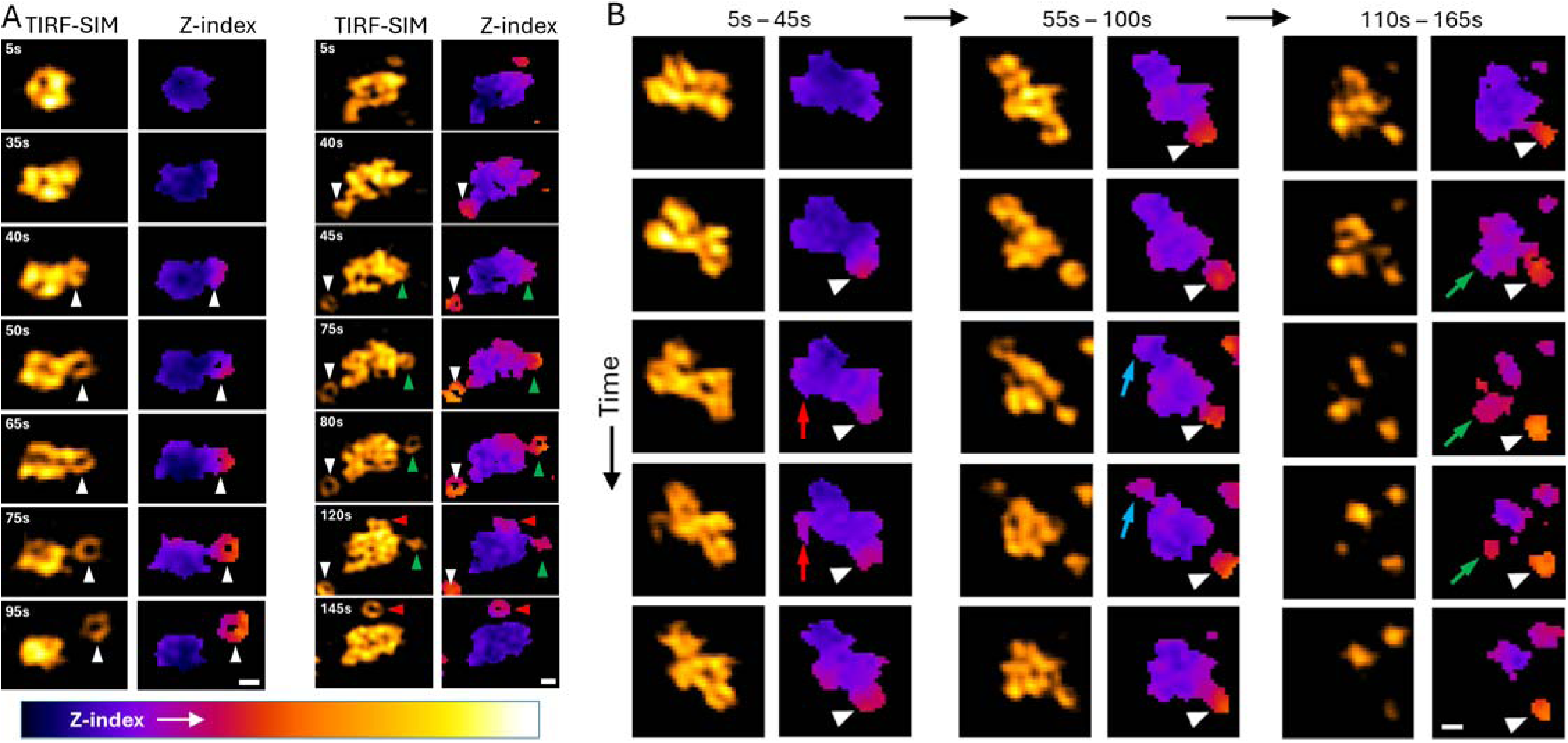
vaTIRF-SIM resolves distinct plaque-associated endocytic events with divergent axial progression dynamics. **A.** Representative examples of clathrin plaques imaged by vaTIRF-SIM, showing TIRF-SIM reconstructions (left) and corresponding Z-index maps (right) over time. Plaque-associated pits (white arrowheads) initiate at peripheral regions of extended clathrin lattices and mature locally, displaying gradual increases in Z-index over tens of seconds prior to internalization. These events remain spatially coupled to the surrounding plaque throughout their lifetime, consistent with localized pit formation at lattice edges (Movies 2&3). **B.** Example of a single clathrin plaque exhibiting two distinct plaque-associated behaviors over time. White arrowheads mark plaque-associated pits that mature slowly with gradual axial progression, whereas colored arrows (red, cyan, green) highlight discrete plaque subdomains that undergo rapid axial progression and are lost from the plaque footprint on shorter timescales (Movie 4). Time proceeds from top to bottom and from left to right as indicated. Scale bars, 0.2 µm.

To integrate lateral organization with axial information, the diffraction-limited axial map was subsequently masked using the SIM-derived segmentation (Figure 1A, 5). This step preserves nanoscale lateral structure while confining the Z-index to clathrin-containing regions, thereby avoiding spatial averaging across heterogeneous assemblies. In the resulting vaTIRF-SIM representation, warmer colors indicate higher Z-index values corresponding to greater axial displacement from the membrane–glass interface, whereas cooler colors indicate more surface-proximal localization.

A representative vaTIRF-SIM image is shown in Figure 1B, demonstrating the ability of this approach to simultaneously resolve clathrin coat architecture and relative axial progression at the plasma membrane. Extended, flat clathrin lattices (plaques; arrows) are readily distinguished from highly curved, ring-shaped clathrin-coated pits (arrowheads), while the Z-index reveals pronounced axial heterogeneity within crowded membrane regions (Li et al. 2015; Nathan M. Willy, Ferguson, et al. 2021). Notably, multiple pit-like invaginations can be detected within or adjacent to a single plaque, each exhibiting distinct axial behavior—features that are obscured in diffraction-limited or purely two-dimensional imaging. Together, these results establish vaTIRF-SIM as a platform for directly linking nanoscale clathrin coat organization with three-dimensional membrane deformation in real time.

### 2.2 Coordinated lateral growth and axial progression during *de novo* clathrin pit formation

To characterize the relationship between lateral clathrin coat organization and three-dimensional progression during *de novo* pit formation, we analyzed individual endocytic events using vaTIRF-SIM (Figure 2). Representative examples illustrate the stereotyped evolution of de novo pits from initial detection as compact assemblies to larger structures that acquire a characteristic ring-like nanoscale organization—an established hallmark of mature clathrin-coated pits—prior to disappearance (Figure 2A)(Li et al. 2015; Nathan M. Willy, Ferguson, et al. 2021). Throughout this process, TIRF-SIM resolves progressive changes in lateral coat architecture, while the corresponding Z-index maps reveal a concomitant increase in relative axial displacement from the plasma membrane.

For individual pits, clathrin fluorescence intensity increased steadily during assembly and peaked prior to pit disappearance, consistent with coat accumulation followed by vesicle release or loss from the evanescent field (Figure 2A, right). In parallel, the maximum Z-index within each pit increased monotonically from the earliest detectable stages of pit formation, indicating that axial progression accompanies clathrin assembly from the outset. Notably, axial progression continued even after clathrin intensity began to plateau, suggesting that inward membrane deformation and associated structural rearrangements persist beyond the phase of maximal coat accumulation (Ferguson et al. 2017; Akatay et al. 2022).

The corresponding average images, computed from the same lifetime-normalized events, further illustrate the coordinated evolution of pit architecture and axial position (Figure 2C) (Movie 1). Early time points (*t* –*t*) are characterized by compact clathrin assemblies with low Z-index values, indicating surface-proximal localization. At later stages (*t*), the averaged TIRF-SIM images display a pronounced ring-like organization, accompanied by elevated Z-index values that spatially coincide with the evolving coat structure. Line profiles taken through the centroid of the averaged pits reveal progressive broadening and redistribution of both intensity and Z-index, consistent with increasing three-dimensional deformation of the membrane as pit maturation proceeds.

To determine whether this behavior reflects a common maturation trajectory, we quantified a population of 75 *de novo* pits and aligned their lifetimes by interpolating each event from first detection to last detection (0–100% completion). Averaging across this population revealed a highly reproducible temporal relationship between lateral coat assembly and axial progression (Figure 2B). Mean clathrin intensity increased early and then plateaued, while the mean maximum Z-index continued to rise over the normalized lifetime, indicating sustained axial progression during pit internalization.

Together, these data demonstrate that *de novo* clathrin-coated pits undergo coordinated lateral reorganization and axial progression throughout their lifetime. Rather than exhibiting a prolonged flat intermediate followed by a discrete topological transition, *de novo* pits display continuous axial progression that accompanies nanoscale reorganization of the clathrin coat. The observed axial progression and nanoscale reorganization are consistent with early curvature generation that intensifies during pit maturation. These findings are in good agreement with previous works using complementary imaging approaches (Nathan M. Willy, Ferguson, et al. 2021; Akatay et al. 2022), which inferred early and progressive curvature generation during *de novo* pit formation, and vaTIRF-SIM now directly visualizes this coupling in real time at nanoscale resolution.

The enhanced lateral resolution afforded by vaTIRF-SIM also enabled us to detect late-stage reorganization events associated with the splitting of mature *de novo* clathrin-coated pits (Figure 3). Importantly, the early stages of these pits are indistinguishable from those of canonical *de novo* events, exhibiting coordinated increases in AP2 fluorescence intensity and Z-index that reflect lateral coat growth accompanied by axial progression. Upon reaching a mature state, marked by the emergence of a ring-like clathrin organization, these pits undergo a pronounced but transient decrease in coat intensity (arrowheads), followed by separation of the coat into two spatially distinct clathrin coat assemblies. The transient nature of the intensity decrease, together with the persistence of elevated Z-index values, indicates that the observed splitting does not coincide with clathrin coat disappearance but occurs during ongoing late-stage pit maturation.

### 2.3 Plaque-associated endocytic events exhibit distinct axial progression dynamics

Having established that isolated *de novo* clathrin pits exhibit coordinated lateral growth and axial progression during maturation, we next examined how these dynamics manifest within extended clathrin plaques, which represent a distinct architectural and mechanical context for endocytosis (Leyton-Puig et al. 2017). Using vaTIRF-SIM, we followed the nanoscale organization and three-dimensional progression of clathrin assemblies that originate within or adjacent to large, persistent lattices with simultaneous lateral super-resolution and dynamic axial readout (Figure 4).

Within plaques, we consistently observed plaque-associated pits (arrowheads) that initiate at the peripheral regions of extended clathrin lattices and mature locally over extended timescales. As shown in Figure 4A, these events display gradual increases in Z-index and persist for tens of seconds prior to internalization, with axial progression comparable to those of isolated *de novo* pits (Movies 2&3). Throughout their lifetime, these pits remain spatially coupled to the surrounding plaque architecture, indicating that curvature develops locally at lattice edges through sequential, pit-by-pit internalization, rather than through global invagination of the plaque. This behavior is consistent with localized lattice remodeling during pit formation, as envisioned by rupture-and-growth–type mechanisms proposed for plaque-associated endocytosis (den Otter and Briels 2011; Lampe et al. 2016; Nathan M. Willy, Ferguson, et al. 2021).

In addition to these slower plaque-associated pits, vaTIRF-SIM resolves a second class of plaque-associated internalization events, characterized by accelerated axial progression and rapid loss from the plaque footprint. These events appear as discrete plaque subdomains that undergo a sharp increase in Z-index followed by disappearance from the evanescent field on substantially shorter timescales than plaque-associated pits (Fig. 4B) (Movie 4). This behavior closely resembles fast plaque internalization events previously described in adherent cells, in which large clathrin assemblies are internalized through an abrupt actin-dependent membrane deformation rather than classical pit-by-pit budding (Saffarian et al. 2009).

Importantly, vaTIRF-SIM enables direct discrimination of these two plaque-associated endocytic modes within the same lattice, as illustrated in Figure 4B, where slowly maturing plaque-associated pits (arrowheads) and rapidly internalizing plaque subdomains (colored arrows) are observed in parallel (Movie 4). By integrating nanoscale lateral organization with dynamic axial progression, vaTIRF-SIM provides the first direct, live-cell comparison of distinct plaque-associated endocytic pathways within a single cellular context.

## 3 DISCUSSION

In this work, we introduce variable-angle TIRF structured illumination microscopy (vaTIRF-SIM) as an imaging framework that integrates enhanced lateral resolution with sensitivity to axial progression near the plasma membrane. While electron microscopy and correlative approaches have provided detailed snapshots of clathrin coat architecture, they are inherently static and cannot capture the dynamic evolution of endocytic events in living cells. Conversely, axial TIRF-based methods—including variable-angle TIRF, polarized TIRF, and dual-wavelength ratiometric approaches—offer sensitivity to membrane-proximal axial position but remain limited by diffraction-limited lateral resolution, restricting their ability to resolve closely spaced pits, plaque subdomains, and dynamic aggregates. By combining TIRF-SIM with variable-angle excitation, vaTIRF-SIM enables simultaneous visualization of nanoscale clathrin coat organization and relative three-dimensional positioning in living cells, without sacrificing temporal resolution. Axial information is extracted through frame-by-frame, pixel-wise ratiometric analysis, yielding a qualitative Z-index that reports relative axial displacement within the evanescent field. Although this approach does not provide absolute axial calibration, it allows direct comparison of axial behavior across individual endocytic structures within the same cellular context. By integrating lateral nanoscale segmentation with dynamic axial readout, vaTIRF-SIM bridges a long-standing gap between structural and dynamic descriptions of clathrin-mediated endocytosis.

Previous models of clathrin-mediated endocytosis have proposed distinct mechanisms for the formation of isolated pits and plaque-associated pits, including flat-to-curved and constant-curvature assembly for *de novo* pits, as well as edge-associated pit formation and rupture-and-growth mechanisms for plaques(den Otter and Briels 2011; Lampe et al. 2016; Nathan M. Willy, Ferguson, et al. 2021). These models were primarily derived from static ultrastructural observations, diffraction-limited live-cell imaging, and theoretical considerations of lattice mechanics, leaving open how lateral lattice organization and three-dimensional progression are coordinated in real time. By integrating lateral super-resolution with dynamic axial readout, vaTIRF-SIM enables direct visualization of these processes in living cells. Applying this approach to *de novo* clathrin-coated pits reveals that axial progression begins early during pit assembly and continues throughout maturation, rather than being confined to a discrete late stage. Population-averaged analyses show coordinated increases in clathrin intensity and Z-index during early and intermediate stages, indicating that nanoscale reorganization of the coat accompanies progressive membrane deformation from the outset. These observations are consistent with early curvature generation that intensifies during pit maturation and are in good agreement with our previous work using complementary imaging strategies (Nathan M. Willy, Ferguson, et al. 2021). By directly resolving both lateral organization and axial behavior, vaTIRF-SIM provides real-time evidence that *de novo* pit maturation proceeds through continuous, coordinated structural evolution rather than through a prolonged flat intermediate followed by a sudden topological transition.

The splitting of mature de novo clathrin-coated pits revealed by vaTIRF-SIM occurs after pits have undergone coordinated assembly and axial progression, indicating that these events arise during late-stage pit maturation rather than from failed or abortive endocytic attempts. In these events, pits initially assemble and progress axially in a manner indistinguishable from canonical *de novo* pits, reaching a mature state marked by a characteristic ring-like organization. Only at this late stage do we observe a pronounced but transient decrease in AP2 fluorescence, followed by separation of the coat into two spatially distinct assemblies. Importantly, this intensity dip occurs while Z-index values remain elevated, indicating that splitting does not coincide with coat disappearance or reversal of axial progression. Instead, the persistence of axial signal suggests continued membrane deformation during a phase of nanoscale coat reorganization. The transient decrease in fluorescence intensity, coupled with persistently elevated Z-index values, indicates that pit splitting reflects reorganization of the clathrin lattice. Such late-stage plasticity is compatible with prior work implicating regulated clathrin lattice remodeling during endocytosis, including roles for Hsc70 and its cofactor GAK in coat rearrangement and turnover (He et al. 2025), although our data do not directly address the molecular drivers of this process. By directly visualizing pit splitting in real time, vaTIRF-SIM uncovers a dynamic remodeling phase that expands current views of pit maturation beyond monotonic assembly and disassembly.

Our analysis reveals clathrin plaques as multifunctional endocytic platforms capable of supporting multiple internalization pathways simultaneously, rather than as uniform or terminal assemblies. Using vaTIRF-SIM, we directly observe that extended clathrin lattices can give rise to at least two distinct modes of endocytosis within the same plaque and over overlapping time windows. One mode consists of plaque-associated pits that initiate preferentially at lattice peripheries and mature locally through gradual axial progression, with kinetics comparable to isolated *de novo* pits. In parallel, plaques also exhibit rapid internalization events in which discrete subdomains undergo accelerated axial progression and are lost from the evanescent field on much shorter timescales. Importantly, vaTIRF-SIM enables these behaviors to be distinguished in real time based on their nanoscale lateral organization and axial dynamics, even when they occur within the same plaque and are indistinguishable under diffraction-limited imaging.

These observations reconcile previously disparate models of plaque-associated endocytosis by placing them within a single spatiotemporal framework. The slowly maturing, peripheral pits are consistent with rupture-and-growth–type mechanisms, in which localized lattice remodeling enables pit-by-pit budding from an extended coat (den Otter and Briels 2011; Lampe et al. 2016; Nathan M. Willy, Ferguson, et al. 2021). In contrast, the rapid plaque subdomain internalization events closely resemble fast, actin-dependent plaque internalization described previously (Saffarian et al. 2009), in which larger portions of the lattice are internalized collectively rather than through sequential pit formation. Rather than representing competing or mutually exclusive pathways, our data indicate that these modes coexist within the same clathrin architecture and may be differentially engaged depending on local mechanical constraints, cytoskeletal coupling, or cargo demands. By resolving nanoscale organization and axial progression simultaneously, vaTIRF-SIM provides the first direct evidence that clathrin plaques function as dynamic and versatile endocytic platforms, integrating multiple internalization strategies within a single lattice.

Several limitations of the present approach should be noted. The Z-index derived from vaTIRF-SIM reports relative axial displacement based on differential evanescent field penetration and therefore provides qualitative, rather than absolute, information about vertical position or membrane curvature. While this is sufficient to compare axial progression dynamics across endocytic structures within the same cellular context, future extensions could incorporate calibrated axial measurements or correlative electron microscopy to directly link Z-index values to ultrastructural geometry. In addition, combining vaTIRF-SIM with markers of actin dynamics (Boulant et al. 2011), or curvature-generating adaptor proteins (Nathan M. Willy, Colombo, et al. 2021) will be important for dissecting the molecular mechanisms that underlie the distinct pit- and plaque-associated behaviors described here. Together, such integrations promise to further refine the spatiotemporal framework introduced in this study and extend its applicability to a broader range of membrane remodeling processes.

## 4 METHODS

### 4.1 Cell Lines and Reagents

Genome-edited human breast cancer SUM159 cells expressing AP2–eGFP (with eGFP fused to the C-terminus of the AP2 σ2 subunit (Aguet et al. 2016)) were maintained at 37 °C in a humidified incubator with 5% CO. SUM159 complete growth medium contains of F-12/GlutaMAX (Thermo Fisher Scientific) supplemented with 5% fetal bovine serum (Gibco), 100 U/mL penicillin–streptomycin (Thermo Fisher Scientific), 1 μg/mL hydrocortisone (H-4001; Sigma-Aldrich), 5 μg/mL insulin (Cell Applications), and 10 mM HEPES (pH 7.4). For live-cell imaging, cells were cultured in L-15 medium (Thermo Fisher Scientific) supplemented with 5% fetal bovine serum and 100 U/mL penicillin–streptomycin, which supports cell growth in the absence of CO equilibration.

### 4.2 Total Internal Reflection and Structured illumination microscopy

A detailed schematic of the home-built high–numerical aperture (NA) structured illumination microscopy system has been previously documented (Akatay et al. 2022). Briefly, a 488 nm laser (300 mW; Sapphire LP, Coherent) is directed through an acousto-optic tunable filter (AA Quanta Tech, AOTFnC-400.650-TN) for wavelength selection and power modulation. The beam is then expanded and relayed to a phase-only modulator to generate structured illumination. The phase-only modulator consisted of a polarizing beam splitter, an achromatic half-wave plate (Bolder Vision Optik, BVO AHWP3), and a ferroelectric spatial light modulator (Forth Dimension Displays, QXGA-3DM-STR), as described previously (Guo et al. 2018).

For high NA TIRF-SIM acquisitions, nine grating patterns consisting of three-orientations and three-phases are displayed on the SLM. The resulting diffracted beams passed through an azimuthally patterned achromatic half-wave plate (Azimuthal HWP; Bolder Vision Optik), which contained three pairs of segments with custom-designed fast-axis orientations. This element rotated the linear polarization of the diffracted light to the desired s-polarization (Guo et al. 2018). A spatial mask was then used to block unwanted diffraction orders, allowing only the ±1 orders to pass. These beams were subsequently relayed to the back focal plane of a high numerical aperture (NA) objective (Olympus APO ×100 1.65 OIL HR 0.15) mounted on an inverted Eclipse TI-E microscope (Nikon instruments Inc.)

For each frame two SIM acquisitions are taken at two incidence angles which are both above the critical angle. One angle is at a higher incidence angle (HiTIRF), and the other angle is at a lower incidence angle (LiTIRF) thereby creating a difference in the penetration depth. The evanescent field penetration depths, described the distance at which the excitation intensity decays to 1/e of its surface value, were estimated to be 55 nm and 75 nm, respectively. The incidence angle at the specimen–coverslip interface (V-A Optical Labs, SF-11) was controlled by adjusting the periodicity of the SLM grating patterns. By adjusting the width of the individual grating in the grating pattern produced by the SLM, we were able to make the incident angle larger or smaller. The smaller the width the greater the incident angle of the beam. The fluorescent emission generated by the applied excitation pattern of each phase and orientation is collected by the same objective and focused by a tube lens onto an sCMOS camera (Hamamatsu ORCA-Fusion BT). The acquired nine raw images are reconstructed into a super-resolution image based using high-fidelity structured illumination microscopy algorithm, HiFi-SIM (Wen et al. 2021). The cells were left in the incubator for 24 h to allow complete spreading before starting imaging using 488 nm illumination at 20 msec acquisition time per frame every five seconds until the cells were photobleached.

### 4.3 Analysis of *de novo* clathrin pits

Individual clathrin-coated pits were manually identified and isolated from TIRF-SIM acquisitions. For each pit, a region of interest (ROI) encompassing the full lifetime of the structure was cropped, beginning at the first frame in which the pit became detectable and ending at the final frame prior to disappearance. All subsequent analyses were performed on these lifetime-aligned ROIs using custom scripts in Python and ImageJ.

#### 4.3.1 Population-averaged temporal dynamics

To compare pit dynamics across events of different durations, each cropped time series was temporally normalized to a fixed length of 40 frames by interpolation. The maximum pixel intensity for each frame of the normalized sequence was then measured to generate the intensity trajectory for each pit. This procedure was applied to both TIRF-SIM and z-index images and the population-averaged dynamics were obtained by averaging these trajectories across all pits (Fig. 2B).

#### 4.3.2 Generation of average pit images

To generate the average-pit reconstructions (Fig. 2C), each cropped pit image sequence was first upscaled laterally by a factor of five using nearest-neighbor expansion to improve spatial sampling for alignment. For each frame, the geometric center of the pit was determined using a binary mask of non-zero pixels. Corresponding frames from the normalized image sequences were then overlaid by aligning their geometric centers, and a pixel-wise mean across these corresponding frames was calculated to generate the average pit image sequence.

## Data availability statement

The raw data supporting the conclusions of this article will be made available by the authors, without undue reservation.

## Ethics statement

The manuscript does not contain any data or descriptions pertaining to human or animal subjects.

## Author contributions

CK conceived the study. CT and GL developed the home-built TIRF-SIM setup and performed the TIRF-SIM measurements. AM performed the *de novo* clathrin pit analyses. CK, CT and AM wrote the manuscript.

